# Force profile of the two-handed hardstyle kettlebell swing performed by an RKC-certified instructor

**DOI:** 10.1101/2021.05.13.444085

**Authors:** Neil J. Meigh, Wayne A. Hing, Ben Schram, Justin W.L. Keogh

## Abstract

**Background:** The effects of hardstyle kettlebell training are increasingly cited in strength and conditioning research, yet reference data from a proficient swing is scarce. The aim of this exploratory study was to investigate the force profile of a two-handed hardstyle swing performed by a Russian Kettlebell Challenge (RKC) instructor.

**Methods:** The subject is a 44-year-old male, body mass 75.6 kg, height 173.5 cm, with six years of regular hardstyle training experience. Two-handed hardstyle swings were performed with a series of incremental mass kettlebells (8-68 kg). Ground reaction force (GRFs) was obtained from a floor-mounted force platform. Force-time curves (FTCs), peak force, forward force, rate of force development (RFD) and swing cadence were investigated.

**Results:** Data revealed the FTC of a proficient swing is highly consistent and dominated by a single force peak (mean SD = 47 N), with a profile that remained largely unchanged to 24 kg. Pearson correlation analysis revealed a very strong positive correlation in peak force with kettlebell mass (*r* = 0.95), which increased disproportionately from the lightest to heaviest kettlebells; net peak force increased from 8.36 ± 0.75 N.kg^-1^ (0.85 × BW) to 12.82 ± 0.39 N.kg^-1^ (1.3× BW). There was a strong negative correlation between RFD and kettlebell mass (*r* = 0.82) that decreased from 39.2 N.s^-1^.kg^-1^ to 21.5 N.s^-1^.kg^-1^. There was a very strong positive correlation in forward ground reaction force with kettlebell mass (*r* = 0.99), expressed as a ratio of vertical ground reaction, that increased from 0.092 (9.2%) to 0.205 (20.5%). Swing cadence exceeded 40 swings per minute (SPM) with all kettlebells.

**Conclusion:** Our findings challenge some of the popular beliefs of the hardstyle kettlebell swing. Consistent with hardstyle practice, and previous kinematic analysis of expert and novice, force-time curves show a characteristic single large force peak, differentiating passive from active shoulder flexion. Ground reaction force did not increase proportionate to kettlebell mass, with a magnitude of forward force smaller than described in practice. These results could be useful for coaches and trainers wanting to improve athletic performance, and healthcare providers using the kettlebell swing for therapeutic purposes. Findings from this study were used to inform the BELL Trial, a pragmatic controlled trial of kettlebell training with older adults. www.anzctr.org.au ACTRN12619001177145.

## Introduction

Kettlebell training has received increasing interest since the first publications in 2009 ^(1, 2)^. Proponents of kettlebell training claim improvements in muscular strength, cardiovascular endurance, explosive power, weight management/fat loss, flexibility, and superior athleticism ^(3)^. Many of the claimed benefits are believed to be attainable from the hardstyle swing popularised by Pavel Tsatsouline. A hardstyle swing is said to have some kinematic similarities with the barbell deadlift and countermovement vertical jump ^(3)^, but the kettlebells’ shape and offset centre of mass, make the kettlebell swing unique, allowing it to be swung between the legs.

National Strength and Conditioning Association (NSCA) standards and guidelines for strength and conditioning professionals, state that knowledge of proper technique, is a cardinal principle of coaching ^(4)^. There is, however, sparce quantitative data of a proficient swing to better understand what McGill and Marshall called “street wisdom” ^(5)^ thus, confidence is low that the current body of evidence is representative of a proficient swing. If hardstyle technique is essential for achieving the claimed effects, it must be clearly defined, in execution and measurement.

A recent review ^(6)^ identified the two-handed hardstyle swing as the most widely investigated kettlebell technique. More than half of the published studies cited Tsatsouline, however, over 80% of study participants were novices. Among 68 studies, it appeared that only four had been conducted by certified hardstyle instructors ^(7-10)^ and only one ^(7)^ provided data from hardstyle-certified practitioners. While certification is not necessary to achieve proficiency, certification could be linked to increased accuracy and reliability, that technique will be performed and assessed consistently. Much remains unknown about the proficient hardstyle swing, especially how this may change as a function of kettlebell load. Thus, a force profile of the proficient swing is warranted.

There are distinct kinematic differences between novice and expert performing a hardstyle swing. A kinematic analysis by Back and colleagues ^(7)^ showed that experts used 20° more hip flexion to perform a ‘hip hinge’, and 19° less shoulder range; the kettlebell being swung upward rather than lifted. A sequence of movements at the hips, pelvis, and shoulders during the upswing and downswing phases of a swing cycle was reported, with the sequence in both phases reversed between expert and novice. During the upswing, experts lead with the hips, followed by the shoulders, whereas novices lead with the shoulders with the hips following. During the downswing, experts allowed the kettlebell to drop before flexing at the hips, while novices flexed at the hips first, with the shoulders following. At the top of the kettlebells’ arc of motion, the experts stood upright in terminal hip and knee extension, but the novices did not. A significant difference in angular velocity at the hips, of 223.8°/s between expert (635.5°/s) and novice (411.7°/s), highlights the ballistic nature of a proficient hardstyle swing. As force is a product of mass and acceleration, differences in ground reaction force (GRF) between expert and novice are expected.

Lake and Lauder ^(11)^ were the first to quantify the mechanical demands of a two-handed hardstyle swing. With a 24 kg kettlebell, peak force was reported to be 19.6 (1.4) N.kg^-1^, impulse 2.5 (0.3) N.kg^-1^.s, and peak power 28.6 (6.6) W.kg^-1^. Horizontal forward force was subsequently reported to be almost 30% of vertical force ^(12)^. Contrary to typical hardstyle practice, the start position was described as “standing still with the kettlebell held in both hands at arm’s length in the ‘finished deadlift’ position, with the kettlebell lightly touching the upper thighs”. The impact of analysing a swing cycle starting from an upright standing position, as opposed to the back or bottom position with the kettlebell between and behind the legs, may be a potential reason that force appears to be *decreasing* throughout the ‘propulsive’ phase.

McGill and Marshall ^(5)^ presented sEMG data of Pavel Tsatsouline performing one and two-handed swings with 32 kg. These data are of interest, but sEMG alone has limited value to practitioners wanting to improve their coaching of the hardstyle swing. With this exception, the study by Back and colleagues ^(7)^ remains the only observation of a hardstyle swing performed by a certified instructor, and the only report of proficient swing kinematics. Prescribing an exercise in the absence of reference standards is challenging for coaches and healthcare providers. This is especially true with athletes, for whom improvements in physical performance is critical, and for higher-risk populations with chronic health conditions. Typical resistance training guidelines for intensity, sets, and repetitions, are not used in hardstyle practice, thus reference standards will help to improve the accuracy and reliability of further research using hardstyle techniques and training protocols.

The aim of this exploratory study was to investigate the force profile of the two-handed hardstyle swing in a certified instructor across a range of kettlebell loads. A representative FTC, peak ground reaction force, rate of force development (RFD) and swing cadence are reported. Results were used to inform the BELL trial (www.anzctr.org.au ACTRN12619001177145).

## Materials & Methods

### Subject

The subject is a 44-year-old male, body mass = 75.6 kg, height = 173.5 cm. He had been regularly training with kettlebells for six years since gaining RKC instructor certification in 2013, including a period of 20 months running group kettlebell classes from a Physiotherapy practice six days a week. The subject was free from injury with no health or medical conditions which would have influenced task performance. Consent was given for the data to be used for scholarly submission, with ethical approval for this study granted by Bond University Human Research Ethics Committee (NM03279).

### Protocol

Data were collected from the University biomechanics laboratory in a single session. The subject performed two-handed kettlebell swings to chest-height on a floor-mounted force flatform (AMTI, Watertown, NY, USA) recording GRF at 1000 Hz using NetForce software (AMTI, USA). Subject body mass was captured by the force plate from a period of quiet standing. Tri-plantar force variables were obtained from the floor-mounted force platform. The variables of interest were peak GRF, forward force, dynamic RFD, and swing cadence. The subject performed a single set of 12 repetitions with each kettlebell, with the middle ten repetitions used for analysis. A custom program (Microsoft Excel, Version 2012) was used to calculate peak force during each swing cycle, with values manually assessed and verified against the corresponding FTC. To obtain net peak force, system weight (body mass + kettlebell mass) was subtracted from the square root of squared and summed data:

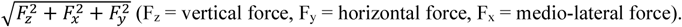

The back or bottom position of the swing was used as the start of each swing cycle. Dynamic RFD (N.s^-1^) during hip extension (propulsion) was calculated as the change in GRF during Phase one, divided by elapsed time and normalised to body mass (N. s^-1^.kg.^-1^), and reported as the mean of ten swings. Cadence in swings per minute (SPM) was calculated from the average time between the peak force during hip extension in each swing cycle – 60s / (Time_peak10_ -Time_peak1_ / 9). Rate of perceived exertion (RPE) was captured for the lightest (8 kg) and heaviest (68 kg) kettlebells. Peak force is reported as resultant force unless stated otherwise.

### Procedure

Swings were performed as described in the RKC Instructor manual ^(13)^. Data from our laboratory indicated that different shod conditions (flat canvas shoes, trainers/sneakers, Oxford lace-up shoes, barefoot), were unlikely to alter the outcome of the study (Supplementary file A). Swings were thus performed barefoot, as recommended. The subject performed a brief self-prescribed mobility drill, with the lighter kettlebells serving as a warm-up for the heavier sets. All sets began and ended with the kettlebell in a dead-start position on the ground. The feet were placed comfortably in the middle of the force platform, at a distance roughly equal to the length of the foot behind the kettlebell, as previously described ^(8)^. Swings were performed with kettlebells from 8 kg to 68 kg. Kettlebell mass increased in increments of 2 kg from 8 kg to 24 kg, then in increments of 4 kg from 24 kg to 48 kg, finishing with 56 kg and 68 kg kettlebells. Kettlebells up to 32 kg were Force USA competition kettlebells of standardised dimensions. Kettlebells 36-68 kg were the ‘traditional’ shape, from Force USA (36-40 kg), Aussie Strength (44-48 kg) and Rogue (56-68 kg). The subject was sufficiently rested between sets.

### The hardstyle swing

The two-handed hardstyle swing is illustrated in Figure 1, consistent with previous studies ^(14) (8)^. Positions in the swing cycle are the ‘start’ (Fig. 1 A), ‘mid-swing’ (E) and ‘end’ (I). The up- and down-swing portions of a swing cycle each have two phases. The first phase of the upswing (propulsion) is described by its primary movement; hip extension. The second phase is the *float* in which the shoulders passively flex. The first phase of the downswing, involving passive extension of the shoulders, is the *drop*. The final phase in the cycle, described as braking or deceleration, sees the kettlebell return to its’ start position. A ballistic movement, resulting in periods of float and drop, distinguishes the hardstyle swing from the double knee bend swing of kettlebell sport.

**Figure 1.**
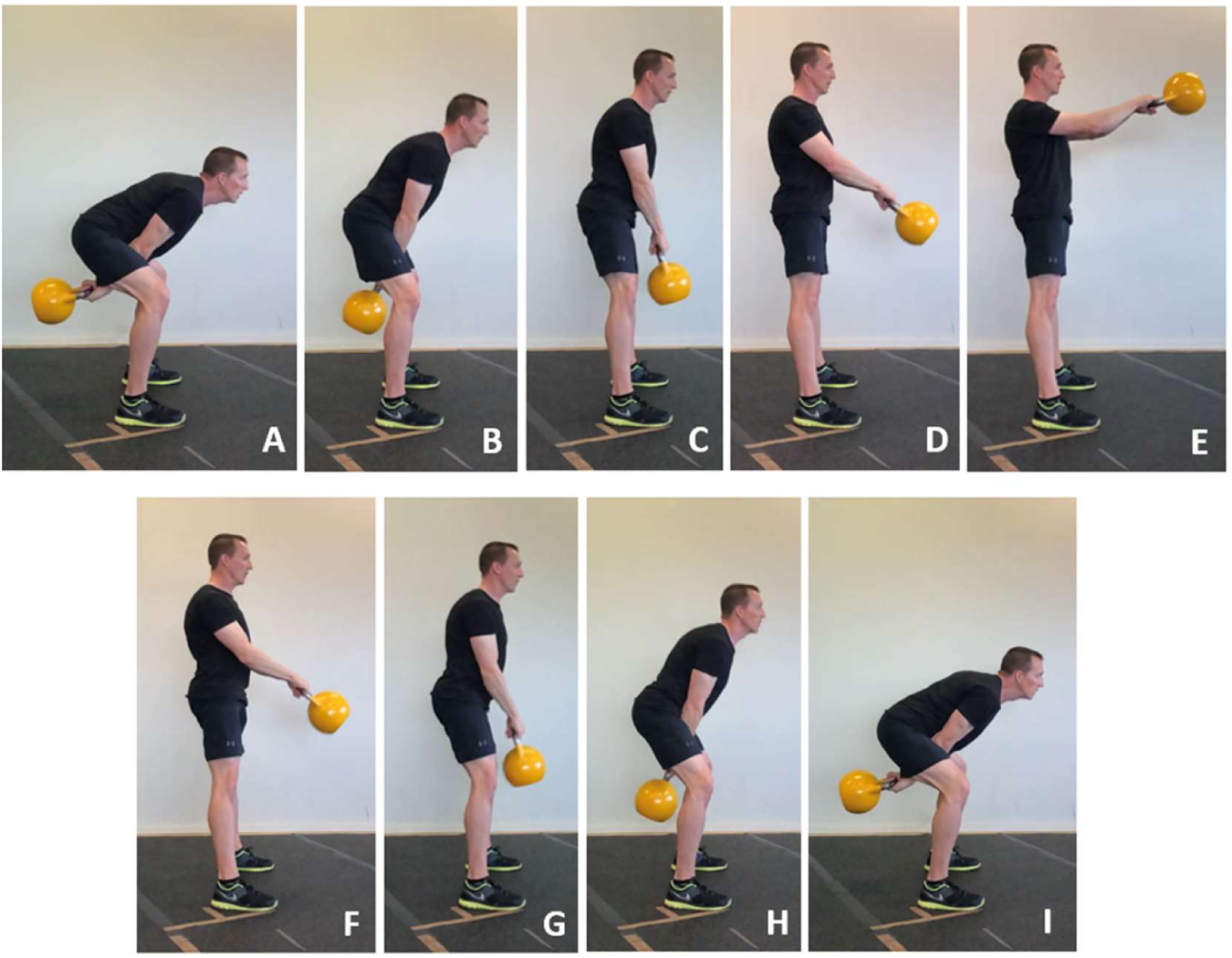
A single two-handed hardstyle swing cycle. A) The Start position of a swing cycle. A-C) Phase 1 - upward *propulsion* of the kettlebell, D-E): Phase 2 - the *float* - passive shoulder flexion with hips and knees in terminal extension, E) *Mid-swing* (top of the swing). E-F): Phase 3 – the *drop* - passive extension of the shoulders with hips and knees in terminal extension, G-I): Phase 4 - *deceleration* (braking) of the kettlebell to the bottom of the swing, I) End of the cycle (bottom of the swing). * subject in the image has Scheuermann’s disease with exaggerated kyphosis.

In the start position, the hips and knees are in terminal flexion (for the exercise), with the kettlebell positioned between the legs and behind the body, mid-forearms in contact with the upper thighs. The trunk is flexed at approximately 45°. Propulsion involves rapid extension of the hips and knees to initiate the kettlebell’s forward and upward trajectory. *“The hips drive explosively from the back swing and then there is a momentary float as the kettlebell reaches the apex of the swing”* ^(13)^. Propulsion ends when the hips and knees reach terminal extension.

During *float*, the shoulders move through passive flexion until the kettlebell reaches its highest vertical displacement, at mid-swing. Throughout *drop*, the hips and knees remain in terminal extension, as the shoulders passively extend, gravity acting on the kettlebell to accelerate it downward. ‘Drop’ ends when the hips and knees begin to flex, the kettlebell at roughly hip height, and upper arms in contact with the ribs. The point at which the person accepts the weight of the kettlebell through the upper limbs, immediately prior to braking, can be described as the ‘catch’. Braking is intended to be borne by the lower limbs, predominantly through eccentric action of the hip extensors. *“at the back of the swing you use the elastic power generated to immediately explode the kettlebell up for another rep. There is no need to swing the kettlebell higher than your chest. The movement of the kettlebell is forward and back, not up and down”* ^(13)^.

Hardstyle swing – RKC standards ^(13)^

1. The back must remain neutral. * At the bottom of the swing, the neck should be slightly extended or neutral.
2. The heels, toes, and balls of the feet must be planted. The knees must track the toes.
3. The working shoulder must be packed [retracted and depressed].
4. During the backswing, the kettlebell handle must pass above knee level.
5. In the bottom position, the working arm must be straight, and elbow locked.
6. There should be no forward movement of the knees or added flexion of the ankles during the upswing.
7. The body should form a straight line at the top of the swing. The hips and knees should be fully extended, and the spine should remain neutral. *
8. Biomechanical breathing should be maintained – exhale when the hips and knees lock out.
9. The abs and glutes should visibly contract at the top of the swing.
10. The kettlebell should float for a moment at the apex of the swing while the hips remain locked out.
11. The hips begin to move back after the upper arm has connected with the ribcage and not before.

### Statistical analyses

Measures of centrality and dispersion are presented as mean ± SD. Effect sizes (ES) were calculated and interpreted using Lenhard and Lenhard ^(15)^ and Magnusson ^(16)^. Effect sizes were quantified as trivial, small, moderate, large, very large, and extremely large where ES < 0.20, 0.20-0.59, 0.60-1.19, 1.20-1.99, 2.0-3.99 and 4.0 respectively ^(17)^. Probability of superiority has been used to illustrate the Cohen’s d effect size, representing the chance that a person from group A will have a higher score than a person picked at random from group B. Linear regression was used to calculate the regression coefficients between the independent variable load, and dependent variables, net peak force, rate of force development, and forward force. Correlations were investigated using Pearson product-moment correlation coefficient, with preliminary analyses performed to ensure no violation of the assumptions of normality, linearity, or homoscedasticity. Data were analysed and linear regression calculated in SPSS (version 26.0; SPSS Inc., Chicago, IL, USA).

## Results

Force-time curves from swings with 8 kg, 16 kg, 28 kg, and 68 kg are presented in Fig. 2. Profiles from all kettlebells are included in Supplementary file B. The FTC of a proficient hardstyle swing cycle is characterised by a tall, single narrow force peak, closely followed by a second distinct force peak of smaller magnitude. The characteristic profile remained consistent from 8-24 kg. It was apparent from the FTC of a 28 kg swing (Fig. 2 C), that the subject was unable to maintain the same duration of float and drop, evident by the progressive merging of force peaks. A visual change to the braking phase duration, was visually evident in the FTC of the 20 kg kettlebell swing (Supp. file B). With cadence remaining relatively unchanged, the duration of the propulsive phase increased with kettlebell mass, with the duration of float and drop phases decreasing. The braking phase, shown as a relatively horizontal force approximating body mass during swings with a 16 kg kettlebell (Fig. 3 A), was almost completely absent in the FTC of a swing with 68 kg (Fig. 3 B). Within-set variability between swing cycles remained small, regardless of kettlebell mass and change in FTC.

**Figure 2.**
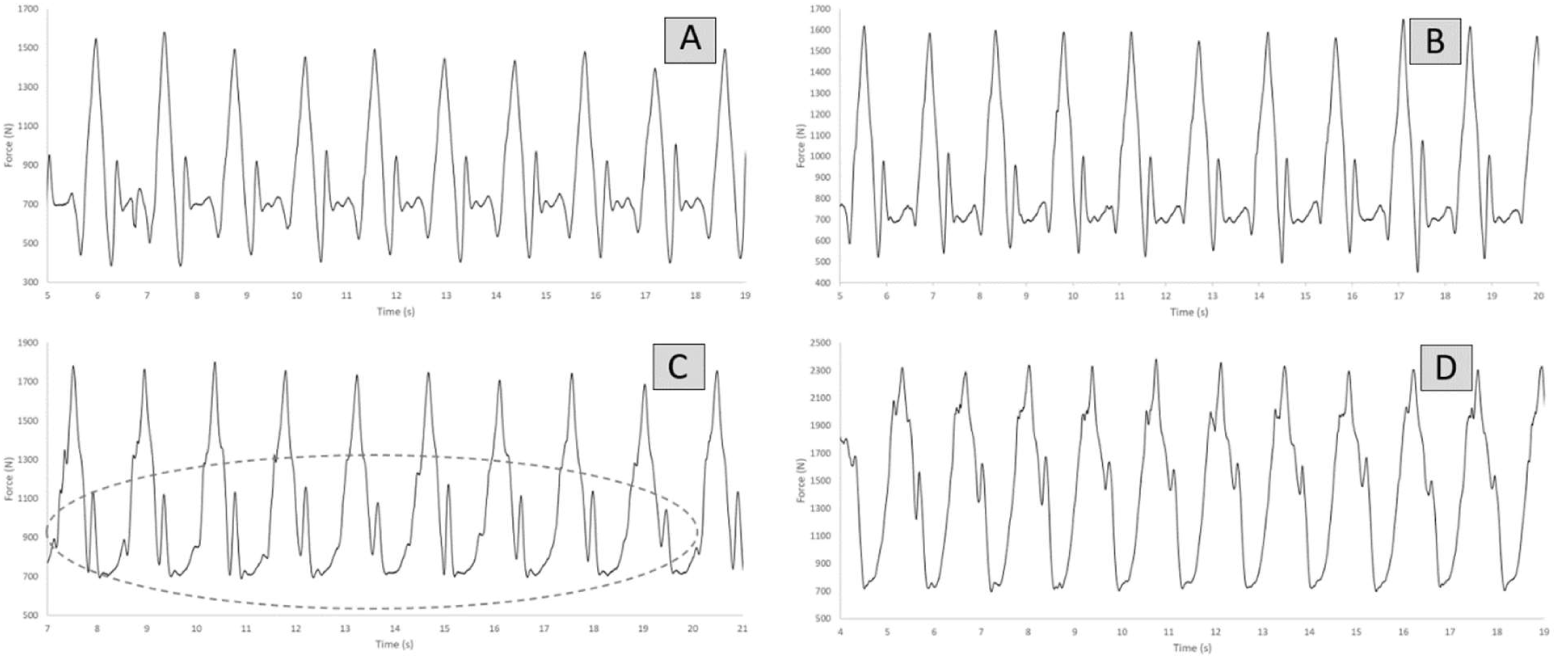
Force-time profiles with (A) 8 kg, (B) 16 kg, (C) 28 kg, (D) 68 kg.

**Figure 3.**
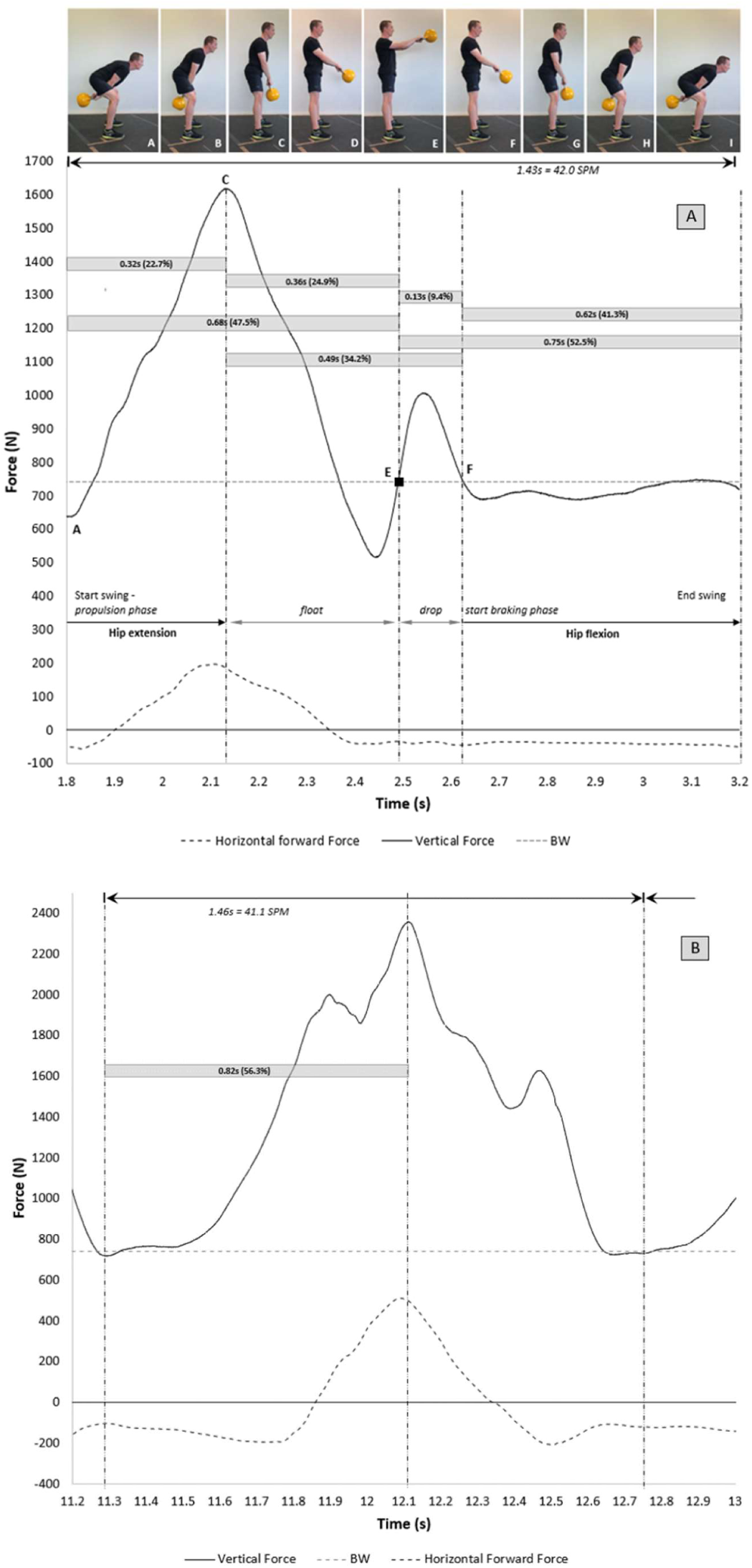
Representative force-time curves of a single two-handed hardstyle swing performed with a 16 kg (A) and 68 kg kettlebell (B). Time and duration of each phase within the 16 kg swing cycle (propulsion, float, drop, braking) have been approximated from the corresponding sequence of images and kinematic sequence of movements. Force = system weight (body mass (741 N) + kettlebell mass (157 N (16 kg), 667 N (68 kg)).

A representative FTC of a single swing cycle with a 16 kg kettlebell is presented in Fig. 3 A. Phase location and durations have been approximated. *Start* and *mid-swing* positions can be reached using a metronome at 80 beats per minute (cadence, 40 swings per minute (SPM)), suggesting that the upward and downward portions of the cycle using light to moderate mass kettlebells are approximately equal (47.5% and 52.5% respectively; Fig. 3 A). Phase 2 & 3 (*float* and *drop*) accounted for approximately one-third of the swing cycle with a 16 kg kettlebell. Peak ground reaction force (1605.9 ± 29.7) was 2.17× BW during the propulsion phase of the upswing, and less than body mass (741 N) for most of the braking phase. A consistent FTC with light to moderate kettlebells (8-24 kg) suggest negligible change in technique. Kettlebells above 24 kg however, influenced swing performance, such that the duration of the float and drop phases progressively diminished.

Figure 3 B shows the considerably altered FTC with the heaviest kettlebell (68 kg). The elapsed time difference in the presented swing cycles is 0.03s (Fig. 3 A = 1.4s, Fig. 3 B = 1.46s). Time to peak force (propulsion) with the 68 kg kettlebell however was 0.82s; >2.5× longer than propulsion with the 16 kg kettlebell. This increased the proportion of the propulsive phase in the swing cycle from 22.7% with 16 kg to 56.3% with 68 kg. A ‘double-peak’ appears to correspond with an accessory effort to move the kettlebell vertically. Movement strategies observed in novices, include active shoulder flexion to ‘lift’ the kettlebell, and excessive extension of the trunk ^(7)^. The braking phase illustrated in the FTC with 16kg (Fig. 3 A), where GRF remains relatively constant, is almost entirely absent with the 68 kg kettlebell (Fig. 3 B). Horizontal forward force expressed relative to vertical force, increased from 0.11 (11%) with 16 kg to 0.21 (21%) with 68 kg.

Figure 4 A-E shows peak force, RFD, forward force, and swing cadence for all kettlebells. There was a very strong positive correlation (*r* = 0.95) in net peak force with kettlebell mass, increasing from 631.8 ± 56.4 N with the 10 kg kettlebell, to 968.4 ± 29.1 N with the 68 kg kettlebell (Fig. 4 A). Net force normalised to body mass increased from 8.36 ± 0.75 N.kg^-1^ to 12.82 ± 0.39 N.kg^-1^; a 1.5× increase in net force, corresponding with an 8.5× increase in kettlebell mass. Normalised net peak force increased from 0.85× BW to 1.3× BW. Subject reported RPE ^(18)^ increased from ‘very, very easy’ (1/10) with 8 kg, to ‘very hard’ (7/10) with 68 kg. There was small within-set variability (SD) in peak force, ranging from 29.1 N with the 68 kg kettlebell, to 76.7 N with the 14 kg (Fib. 4, B), with a mean SD of 47.0 N. Variability in RFD is larger with the 28 kg, 32 kg and 36 kg kettlebells due the change in FTC making the start of hip extension less distinct. There was a strong negative correlation (*r* = 0.82) in RFD with kettlebell mass, decreasing from 39.2 N.s^-1^.kg with the 8 kg kettlebell, to 21.5 N.s^-1^.kg with the 40 kg kettlebell (Fig. 4 C). There was a very strong positive correlation (*r* = 0.99) in forward force with kettlebell mass, from 0.092 (9.2%) to 0.205 (20.5%) (Fig. 4 D). There was only a weak positive correlation (*r* = 0.41) in swing cadence with kettlebell mass, increasing from 41.1 SPM with 20 kg to 44.0 SPM with 68 kg. Cadence for all swing sets was higher than the subject’s self-reported usual training cadence (40 SPM).

**Figure 4.**
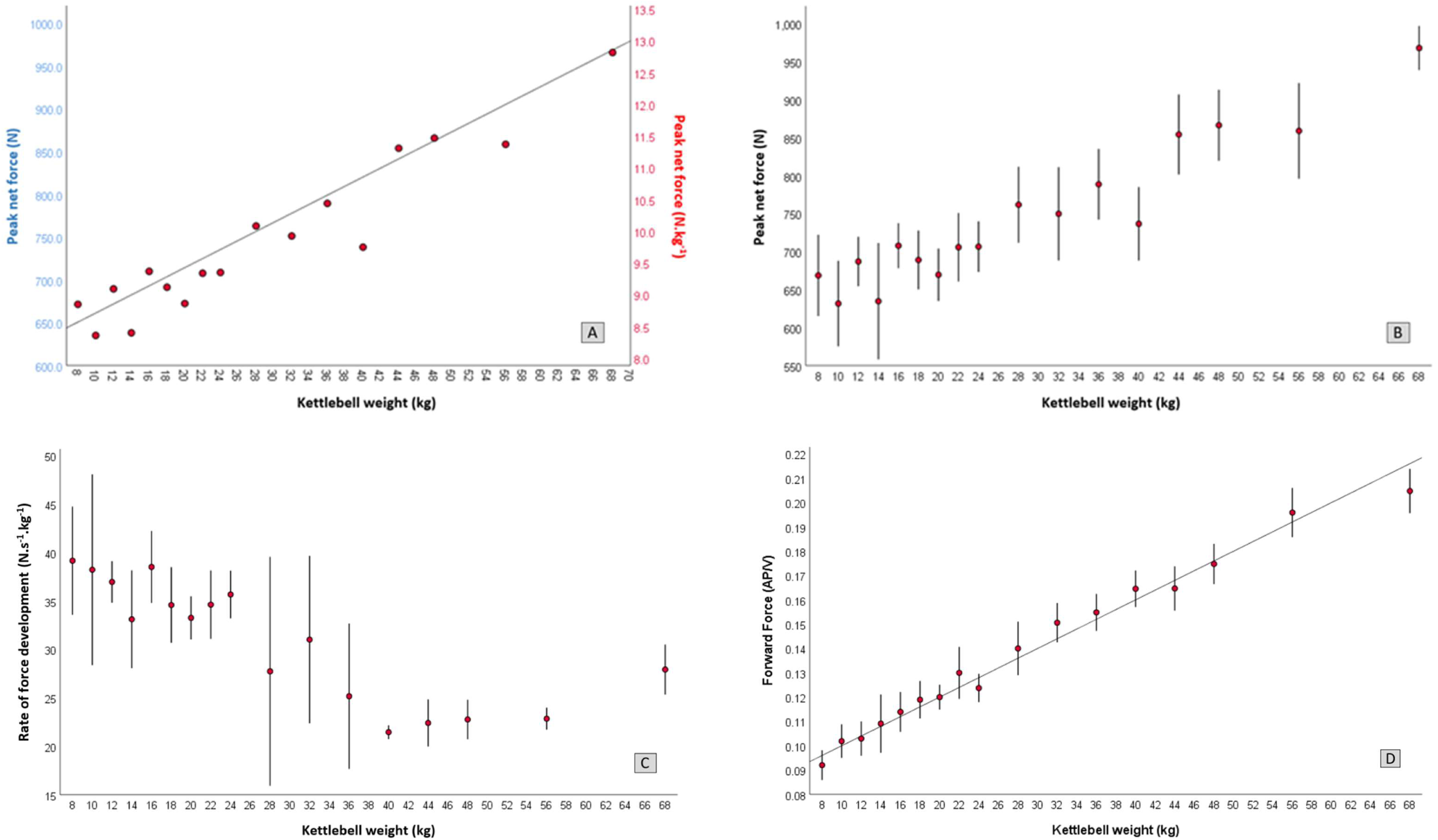
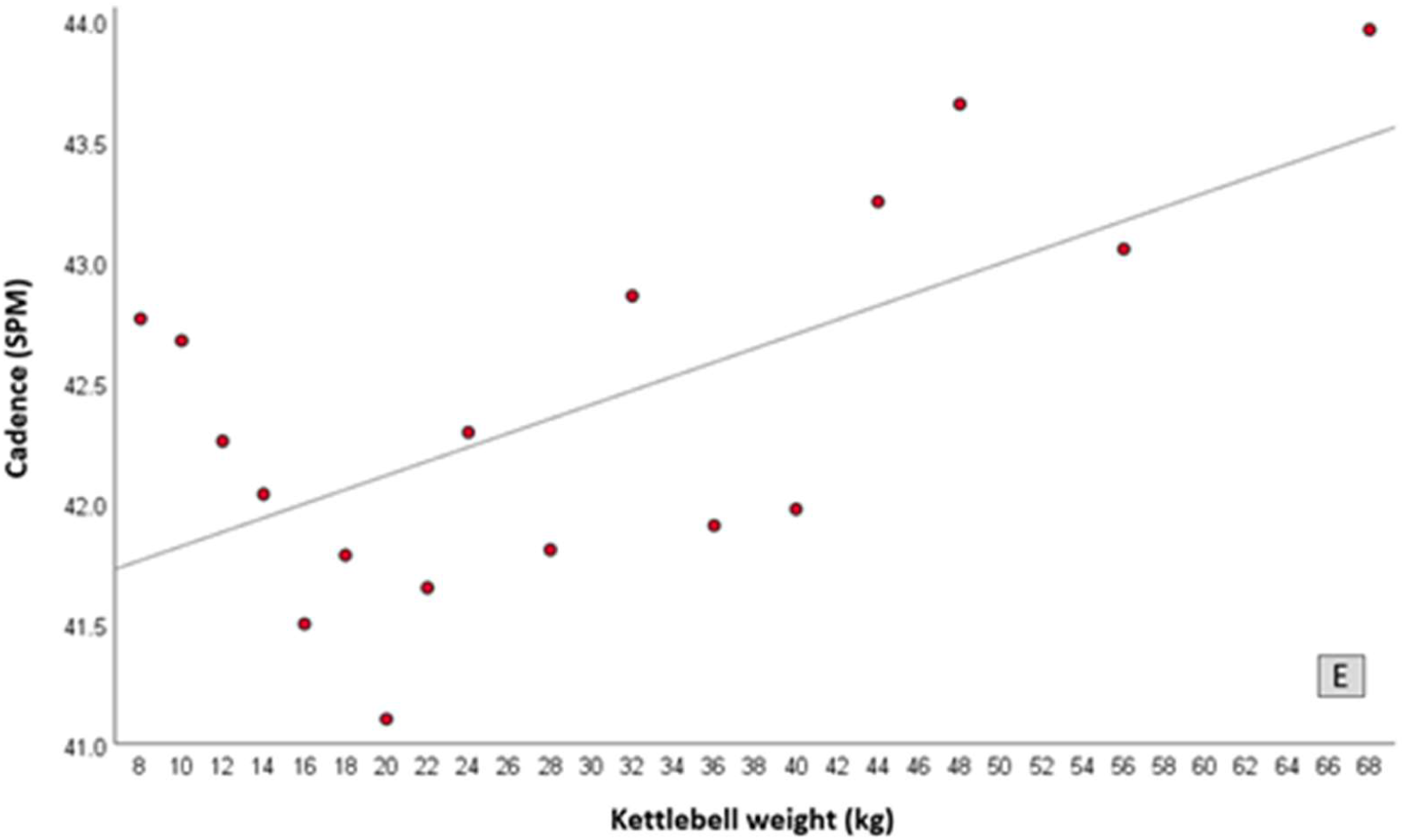
Change in force variables and swing cadence with increasing kettlebell mass. A: net peak force, B: variability in net peak force, C: rate of force development, D: ratio of forward horizontal force to vertical force (A = anterior, V = vertical), E: swing cadence (SPM).

Force variables and cadence for 8-32 kg kettlebells are presented in Figure 5 and Table 1. Linear regression equations are presented in Table 2.

**Table 1.**
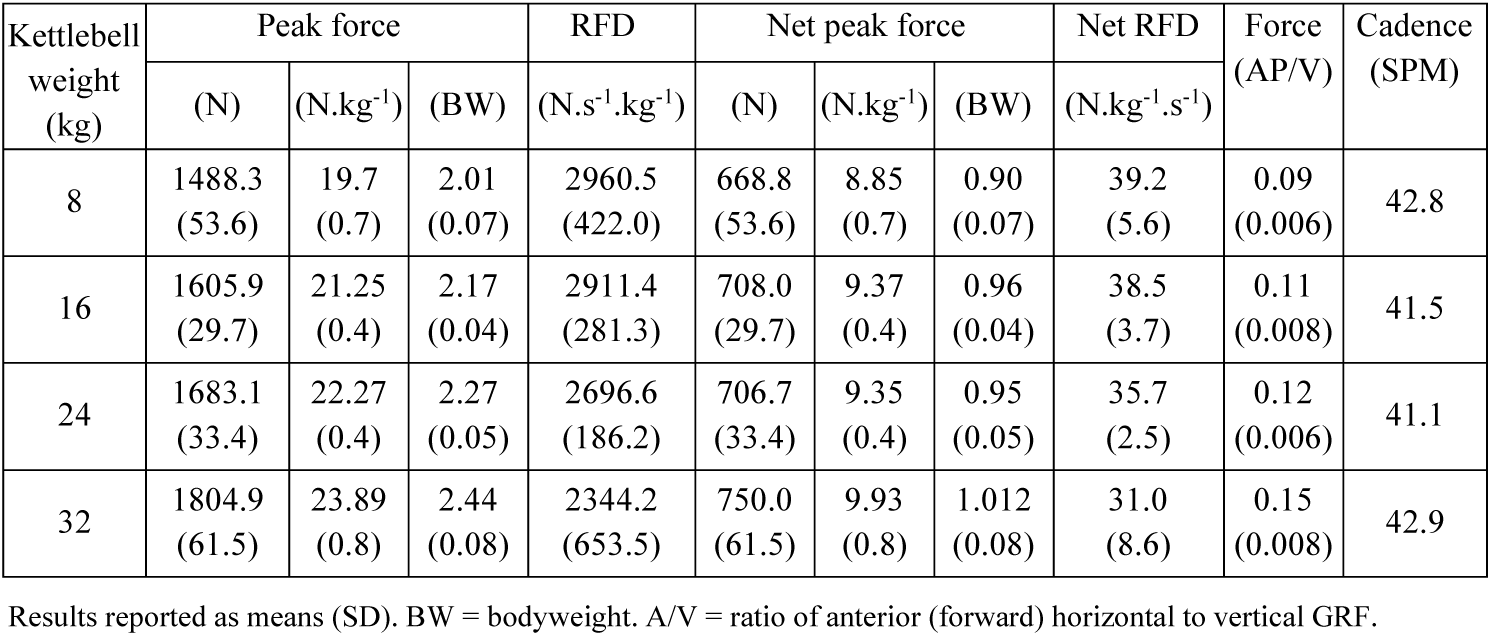
Force variables and cadence with increasing kettlebell mass. Most frequently used and commercially available kettlebells (8-32 kg).

| Kettlebell weight (kg) | Peak force |  |  | RFD | Net peak force |  |  | Net RFD | Force (AP/V) | Cadence (SPM) |
| --- | --- | --- | --- | --- | --- | --- | --- | --- | --- | --- |
|  | (N) | (N.kg<sup>-1</sup>) | (BW) | (N.s<sup>-1</sup>.kg<sup>-1</sup>) | (N) | (N.kg<sup>-1</sup>) | (BW) | (N.kg<sup>-1</sup>.s<sup>-1</sup>) |  |  |
| 8 | 1488.3<br>(53.6) | 19.7<br>(0.7) | 2.01<br>(0.07) | 2960.5<br>(422.0) | 668.8<br>(53.6) | 8.85<br>(0.7) | 0.90<br>(0.07) | 39.2<br>(5.6) | 0.09<br>(0.006) | 42.8 |
| 16 | 1605.9<br>(29.7) | 21.25<br>(0.4) | 2.17<br>(0.04) | 2911.4<br>(281.3) | 708.0<br>(29.7) | 9.37<br>(0.4) | 0.96<br>(0.04) | 38.5<br>(3.7) | 0.11<br>(0.008) | 41.5 |
| 24 | 1683.1<br>(33.4) | 22.27<br>(0.4) | 2.27<br>(0.05) | 2696.6<br>(186.2) | 706.7<br>(33.4) | 9.35<br>(0.4) | 0.95<br>(0.05) | 35.7<br>(2.5) | 0.12<br>(0.006) | 41.1 |
| 32 | 1804.9<br>(61.5) | 23.89<br>(0.8) | 2.44<br>(0.08) | 2344.2<br>(653.5) | 750.0<br>(61.5) | 9.93<br>(0.8) | 1.012<br>(0.08) | 31.0<br>(8.6) | 0.15<br>(0.008) | 42.9 |

Results reported as means (SD). BW = bodyweight. A/V = ratio of anterior (forward) horizontal to vertical GRF.

**Table 2.**
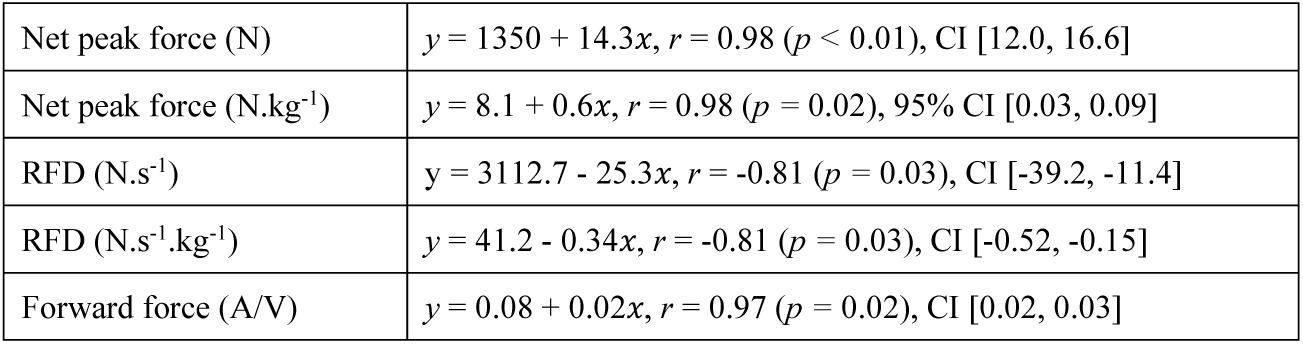
Linear regression equations for net peak GRF, RFD, and forward force forward force relative to vertical. force. *x* = kettlebell mass in kg.

**Figure 5.**
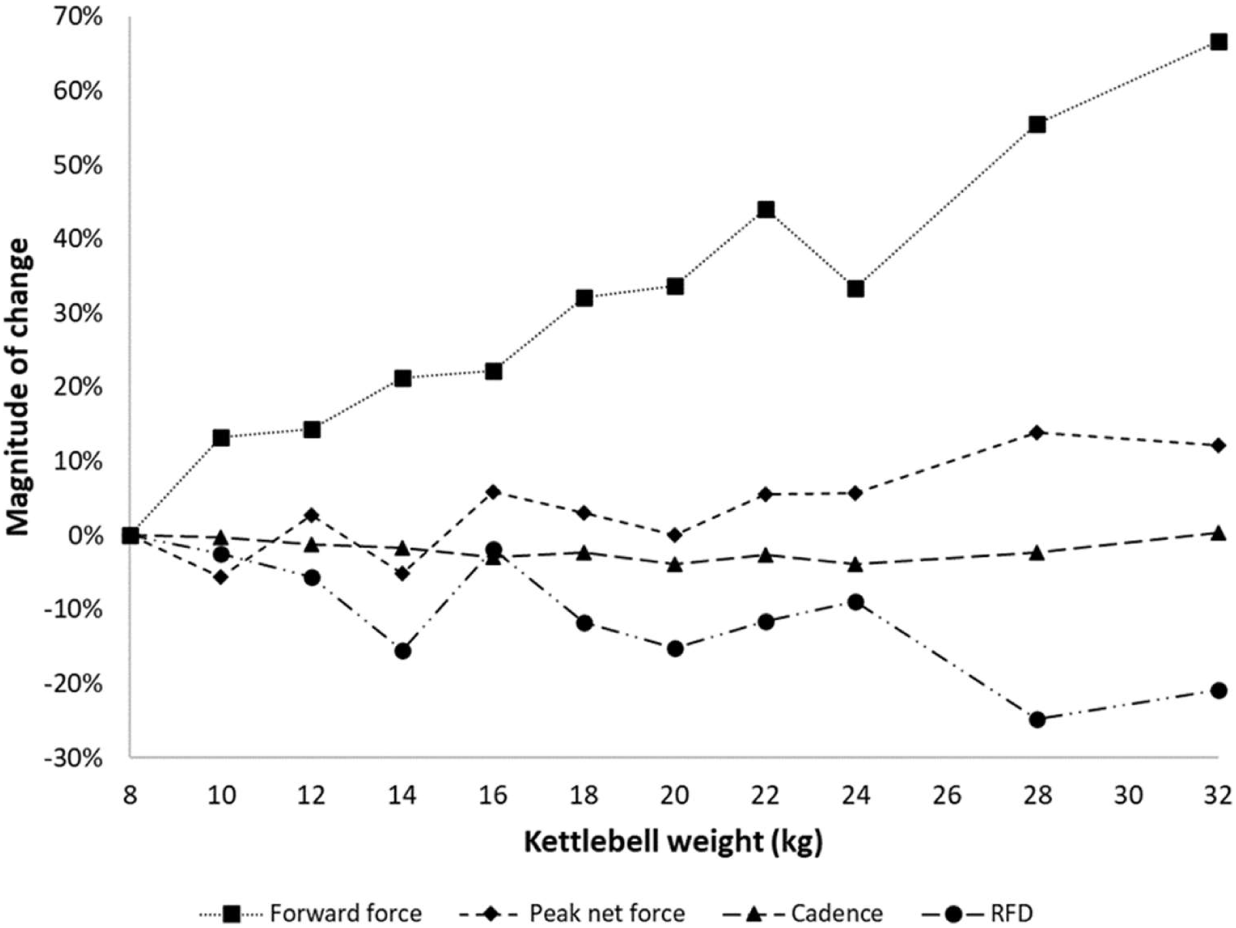
Magnitude of change with increasing kettlebell mass, from 8 kg to 32 kg.

There were very strong positive correlations between kettlebell mass and peak force, and kettlebell mass and forward force. For each 1 kg increase in kettlebell mass, net peak force increased by 12 to 16.5 N.kg^-1^ and forward force by 2 to 3%. There was a strong negative correlation between kettlebell mass and RFD; for each 1 kg increase in kettlebell mass, RFD decreased by 11 to 39 N.s^-1^. Swing cadence remained relatively stable at 42.0 ± 0.6 SPM.

## Discussion

Reliable kinetic reference data enables coaches and healthcare providers to make informed decisions about the potential benefits, or risks, of an exercise. These data can also provide valuable insights which influence how exercises are coached and tested. A properly designed program using resistance equipment, includes multi-joint exercises, such as the kettlebell swing, is individualised, periodised, progressive, and includes appropriate technique instruction ^(19)^. A recent review ^(6)^ was unable to identify data for any of these program variables, which had either been derived from proficient swings, or were appropriate for healthcare providers. Results from the present case study provide some insight, for coaches and healthcare providers, to make more informed decisions about how to use the kettlebell swing for performance enhancement or the prescription of therapeutic exercise.

### Peak force

The magnitude of change in peak force with increasing kettlebell mass was surprisingly small. Quadrupling the kettlebell mass, from 8 to 32 kg, increased peak force by only 81.2 N. Net peak force increased by less than 30% (190 N) between the 8 and 56 kg swings. *“If your goal is more force production, swing a heavier kettlebell”* ^(13)^ appears to be a somewhat misguided instruction. A strong correlation between GRF and kettlebell mass, all the way up to 68 kg, was also surprising. The rate of increase in net force had been expected to slow considerably with heavier kettlebells e.g., >32kg, but this was not the case.

Lake and Lauder ^(11)^ reported peak force during swings with 16 kg, 24 kg, and 32 kg as 18.8 (0.5), 19.6 (1.4) and 21.5 (1.4) N.kg^-1^, respectively. Although absolute difference in mean values between the instructor in the present study, and those reported by Lake were small (range = 2.39 to 2.67 N.kg^-1^), the ES were large to very large (16 kg: δ = 4.9, 24 kg: δ = 1.91, 32 kg: δ = 1.71) with 88.7% to 100% probability of superiority in favour of the instructor. These data suggest that proficiency (technique) is likely to alter ground reaction with large effect.

With kettlebells from 10-20% BW, Levine and colleagues ^(20)^ reported peak GRF from 1.53 (0.2) to 1.67 (1.7) BW. The ES with comparable kettlebells in the present study, was very large, ranging from δ = 2.09 (8 kg, 10% BW) to δ = 2.72 (12 kg, 15% BW), with a mean probability of superiority >95%. Bullock and colleagues ^(21)^ reported peak vertical force of 0.98 (0.1) BW, however data was not reported as net of system weight. In the present study, vertical GRF was 2.15 (0.1) BW with the same kettlebell (20 kg). The ES difference between studies is so unreasonably large, as to suggest that the data (or comparison) is unreliable. If the vertical force reported by Bullock and colleagues was indeed net force, there would be a small effect size difference in favour of the novices (12 kg: δ = 0.36, 20 kg: δ = 0.57). While not impossible, the data from Lake and Lauder ^(11)^ suggests this scenario would be unlikely.

Among male kettlebell sport competitors, Ross and colleagues ^(22)^ reported peak GRF during a 24 kg kettlebell snatch as 2.10 (0.31) BW. In the present study, peak force during the 24 kg swing was 2.27 (0.05) BW. As force is a product of mass and acceleration, the small ES in favour of the swing (δ = 0.55) is most likely explained by the pronounced differences in cadence; 13.9 (3.3) snatches overhead per minute vs 42.3 SPM to chest height. These data appear otherwise comparable, suggesting that proficiency is more likely to influence the force profile and its associated effects, than the ‘style’ of kettlebell training (hardstyle vs Sport). This similarity in GRF also underscores the demands of a unilateral kettlebell exercise (swing, clean, snatch). This highlights an opportunity for coaches and healthcare providers to increase the physiological demand of the swing without increasing the kettlebell mass.

### Rate of force development

Rate of force development, claimed to be a defining feature of the hardstyle swing, is essential for sports performance. It is also an important consideration in rehabilitation, return to sport following injury, and critical for trip and falls prevention ^(23)^. In older adults, RFD may be more important than strength, with respect to functional performance and maintaining independence ^(24-26)^. If the intent of a kettlebell swing is to improve RFD, the findings of this study suggest that even moderate-load kettlebells (16-24 kg) may be counterproductive, with the lightest kettlebells (8 kg) producing the highest RFD (38.2 N.s^-1^.kg^-1^). In the present study, swings performed with 40 kg (equal to 50% 10RM), resulted in a 45% reduction in RFD compared to the 8 kg kettlebell.

*“When you cannot maintain your explosiveness any longer, it’s time to quit*.*”* ^(13)^. The subject in the present study maintained desirable form up to 68 kg, yet the FTC and relative “explosiveness” had evidently changed considerably. If explosiveness can be characterized by the RFD and swing cycle phase duration, and maximising explosiveness is the training goal, we propose that merging of the force peaks could be used to determine a maximum training load. In this case, merging of the force peaks was evident with a 28 kg kettlebell. Kettlebells under 25% body mass, appear to be most suited for improving RFD in a two-handed hardstyle swing.

### Exercise prescription and coaching

Tsatsouline ^(3)^ recommends that an average male beginner starts with a 16 kg kettlebell; 32 kg kettlebells being reserved for “advanced men”, stating, *“unless you are a powerlifter or a strongman, you have no business starting with a [24 kg]”*. Tsatsouline suggests that an average woman should start with an 8 kg kettlebell and a strong woman 12 kg, with most women advancing to 16 kg. These starting kettlebells have frequently been used in research, but these are general guidelines for kettlebell training which includes other exercises besides the swing e.g., the military press, goblet squat, snatch, and Turkish get-up. A kettlebell best suited for a swing is unlikely to be the most appropriate for pressing overhead or deadlifting. If the kettlebell swing is to be performed explosively, with the aim to improve muscular power and functional performance, the results of the current study may suggest that slightly lighter loads than recommended by Tsatsouline ^(3)^ might provide the best outcomes for most individuals. However, these recommendations might still need to be changed based on the size and strength of the participant; such loads may be too light for a 110 kg strength athlete but too heavy for a 45 kg septuagenarian with osteoporosis. Choosing the most appropriate kettlebell for a person performing a given kettlebell exercise, should be established at an individual level, with consideration given to factors such as training age, physical capacity, and training goals.

*“The swing is an expression of forward force projection such as found in boxing or martial arts, like a straight punch*.*”* ^(27)^. A hardstyle swing is defined by its dominant movement - hip extension. Instruction to drive the hips forward rapidly and aggressively, has translated to a belief that the dominant ground reaction is also in the forward horizontal direction. These data do not support that inference. Contrary to popular commentary, forward force ranged from 9% to 21%, with the median being just 13%. The difference in magnitude of forward force during swings with a 24 kg kettlebell, between the 30% reported by Lake and Lauder ^(11)^, and the 12% reported in the present study, is most likely explained by the different starting positions.

These data suggest that centrifugal force acting on the kettlebell is the result of a predominant (≈85-95%) vertical ground reaction. Training instruction encouraging a movement pattern consistent with a vertical jumping motion, rather than attempting to project the kettlebell forward, is likely to be more effective. Investigation of the influence of technique proficiency on the hammer throw ^(28)^, shows a shift in centre of mass, significantly alters the pendular arc of motion and resultant throwing distance. With similar observations in elite kettlebell sport athletes ^(29)^, it appears that small changes in technique, are most likely to account for the large differences in GRF observed between expert and novice, highlighting the role and potential value of expert instruction and coaching.

An ideal hardstyle swing, can be visually identified, by observing the person’s body position at the beginning of the float (Fig. 1 D) and the end of the drop (Fig. 1 F); they should look the same with the direction of kettlebell travel not apparent from the body position. Use of slow-motion video analysis to provide feedback is encouraged as an effective teaching strategy ^(30)^. Real-time biofeedback from a force platform could also be a useful tool for coaches and healthcare providers. Previous published FTCs of the hardstyle swing ^(11, 31)^ show a wide multi-peaked force profile, which is inconsistent with the single, narrow force peaks found in the present study. We propose that a multi-peaked FTC is representative of the active shoulder flexion described by Back and colleagues ^(7)^. If phase durations are important and FTC characteristics a reliable indicator of proficiency, an FTC might be helpful in establishing the optimal swing load.

Proficient hardstyle practitioners perform swings at a cadence of around 40 SPM ^(8, 10, 32)^. The subject in the present study also self-reported a usual training cadence of 40 SPM. The slighter higher mean cadence of 42 SPM in the current study is attributed to the test conditions. While cadence may be intentionally increased or decreased, this is reported to feel “unnatural” ^(10)^, requires greater effort ^(32)^, and is not recommended. *“Don’t confuse a quick and explosive hip drive with manic speed. Pulling the kettlebell down and releasing your hips too soon without allowing the bell to float will give the sense that you are increasing speed, but it will not increase power production”* ^(13)^.

To optimise outcomes, prescription of a kettlebell swing should be personalised. Coaches and healthcare providers will need to determine for the individual, if it is necessary or beneficial for kettlebell mass (intensity) to be increased, or whether the same training effect can be achieved with a higher number of repetitions using a lighter kettlebell. Swings allow a large volume of work to be performed in a short period of time. If the performance goal is simply to get ‘Work’ done, heavier kettlebells are an attractive option; what can be achieved in 90 seconds with a 40 kg kettlebell using a 1:1 work:rest ratio of 2×20 reps (1600 kg), would take 9m:30s at the same continuous pace with an 8 kg kettlebell.

The difference in training loading volume between 5 sets of 10 swings with an 8 kg, 16 kg, or 40 kg kettlebell, is a substantial 1,600 kg. The difference in cardiovascular response and effort is also likely to be very high. Results from this study show a disproportionate increase in effort relative to kettlebell mass and the net peak force. Training parameters of exercise, load, sets, repetitions, and work:rest ratio are important variables for prescription, especially for individuals with higher-risk health conditions such as cardiovascular disease. Resistance training improves important metabolic parameters in people living with major non-communicable diseases, including type 2 diabetes, cardiovascular disease and cancer, but data are limited ^(33)^. Further research is warranted to help coaches and healthcare providers determine safe and effective parameters for prescribing kettlebell exercises with at-risk populations.

Increasing kettlebell mass or cadence increases cardiovascular response ^(10)^ however, a slower cadence can also increase effort ^(32)^ as ‘swing’ and ‘drop’ become a ‘lift’ and ‘lower’. A metronome can be used as an external cue to optimise efficiency in the hardstyle swing. Coaches and healthcare providers should also note that the cardiovascular demand of kettlebell swings is greater than walking ^(8)^ and anticipate that heart rate (HR) will increase with continuous swings ^(34)^ potentially to a relatively high percentage of HR_max_. Similar effects may also occur when performing multiple sets of swings with short periods of rest between sets ^(35)^. The magnitude of these cardiovascular responses may also need to be considered, when prescribing kettlebell swing training to different individuals, especially clinical patients with compromised cardiovascular and/or respiratory function.

### Strengths and limitations

The strengths of this study include the use of an RKC-certified kettlebell instructor, and large range in kettlebells. However, as this paper provides data from one certified kettlebell instructor, the results cannot be generalised to all instructors; replication is necessary to increase our confidence in the results. These data are from ground reaction only; concurrent motion capture would elucidate the kinetic and kinematic changes imposed by increasing kettlebell mass. Reliability in calculating RFD from an FTC may also be compromised where the onset of propulsion is not clear, meaning a more reliable standardised measure of calculating RFD is warranted. Ground reaction data from hardstyle swings cannot be generalised to the double knee-bend (kettlebell sport) or overhead (American) swings which are kinematically different ^(36, 37)^.

## Conclusions

The aim of this paper, was to investigate the force profile of a two-handed hardstyle kettlebell swing, performed by an RKC-certified instructor. The force-time profile using light to moderate mass, was characterised by a smooth, single narrow force peak, immediately followed by a second peak of smaller magnitude. Small within- and between-set variability was observed, with cadence no less than 40 swings per minute. Flight time accounted for approximately one third of the swing cycle, for kettlebells up to approximately 25% body mass. With increasing load, propulsion and braking phases increased, evident by progressive merging of the force peaks.

There were very strong positive correlations between kettlebell mass and peak force, and kettlebell mass and forward force, although the magnitude of change was small. Net peak force increased by less than 30% from swings with an 8 kg kettlebell to 56 kg. More research is required to determine the potential benefits of performing swings with very heavy kettlebells.

Median horizontal forward force was less than 15% of vertical force, indicating that the hardstyle swing is an expression of a predominantly vertical ground reaction. Cues which replicate a vertical jumping motion, appear to be the most useful coaching strategy to optimate swing performance. There was a strong negative correlation with rate of force development. Kettlebells under 25% body mass appear to be optimal for developing lower limb power, with the lightest kettlebells resulting in the highest RFD.

Results show that proficiency in hardstyle technique changes ground reaction with large effect. Proficiency should be considered when reporting and interpreting data from novices. Developing a proficient hardstyle swing is likely to be beneficial, where force and other mechanical demands are important. Further research is required to better understand the effects of kettlebell weight on outcomes of interest, and prescription variables within a hardstyle program.

### Plain language summary

Net ground reaction force during two-handed swing with an 8 kg kettlebell was approximately double bodyweight. Around 90% of the ground reaction force during the swing is vertical and its magnitude, relative to bodyweight, appears to be significantly influenced by proficiency. Peak force during a proficient swing with a 24 kg kettlebell is comparable to that of a 24 kg snatch performed by a proficient Kettlebell Sport practitioner. Effort reported during the swing increased disproportionately to the mass of the kettlebell, and the greatest rate of force development is expressed with the lightest kettlebells. The force-time curve of a proficient swing is characterised by a single force peak during the up-swing, whereas a novice swing is characterised by a double peak.

## Acknowledgements

The authors would like to acknowledge Mr Benjamin Hindle for his support in the calculation of rate of force development, and Mrs Evelyne Rathbone for her direction with statistical analysis

## Contributions

NM conducted the study, curated, and analysed the data, interpreted the results, conducted the formal analysis, and wrote the original draft. JK, BS and WH supported with ongoing consultation. JK and WH reviewed and provided revisions to earlier versions of the manuscript. All authors read and approved the final manuscript.

## Ethics declarations

### Ethics approval and consent to participate

Not applicable

### Consent for publication

Not applicable

### Rights and permissions

Open Access This article is distributed under the terms of the Creative Commons Attribution 4.0 International License (http://creativecommons.org/licenses/by/4.0/), which permits unrestricted use, distribution, and reproduction in any medium, provided you give appropriate credit to the original author(s) and the source, provide a link to the Creative Commons license, and indicate if changes were made. The Creative Commons Public Domain Dedication waiver (http://creativecommons.org/publicdomain/zero/1.0/) applies to the data made available in this article, unless otherwise stated.

