## Supplementary figures and images for "Force profile of the two-handed hardstyle kettlebell swing performed by an RKC-certified instructor"

### Supplementary file A

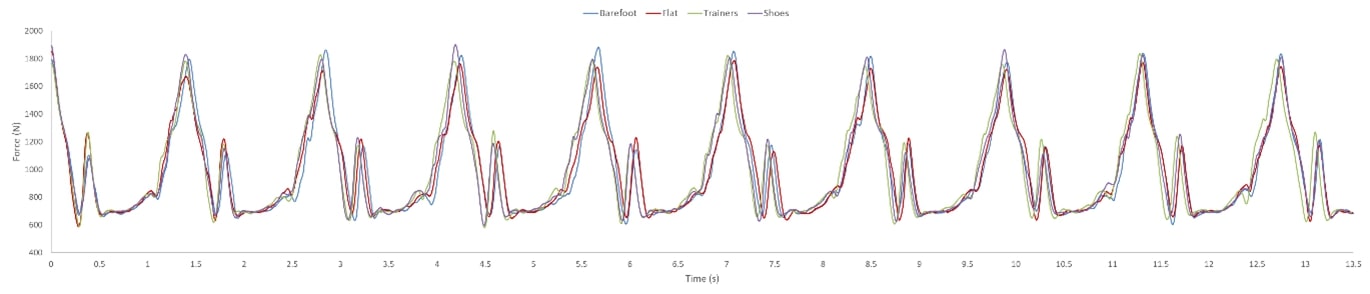

### Supplementary file B

8 kg

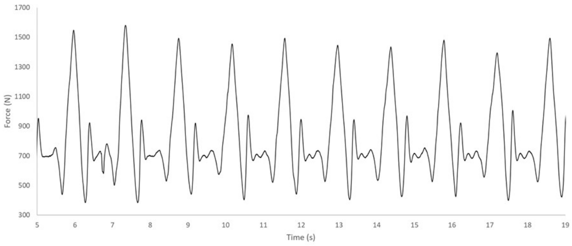


10 kg

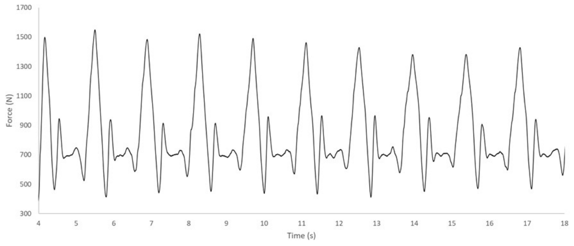


12 kg

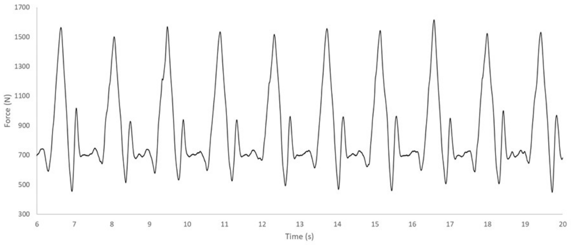


14 kg

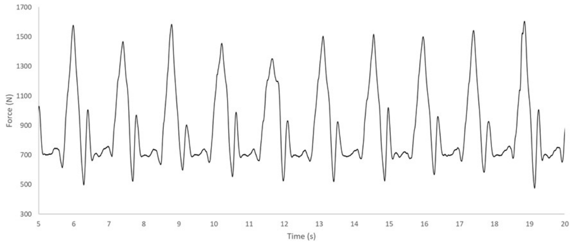


16 kg

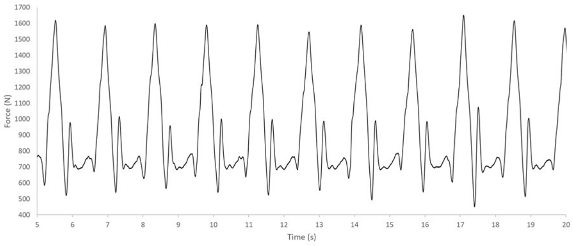


18 kg

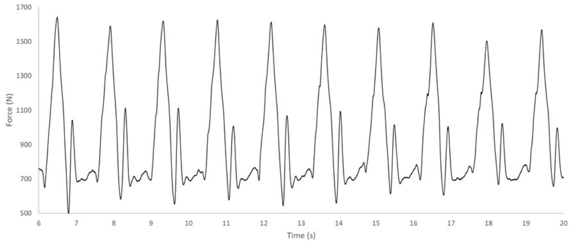


20 kg

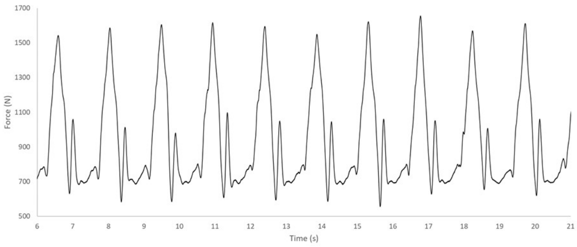


22 kg

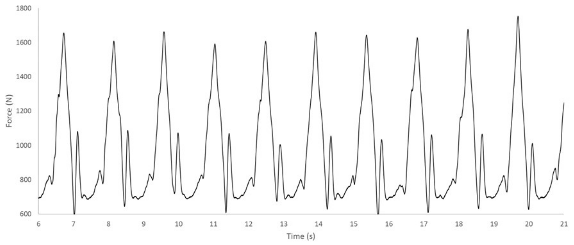


24 kg

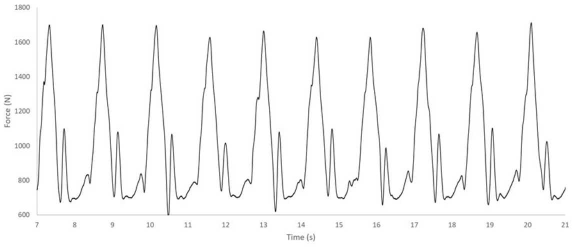


28 kg

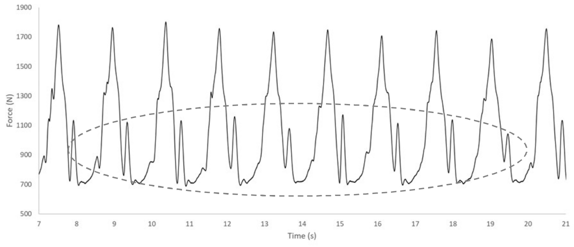


32 kg

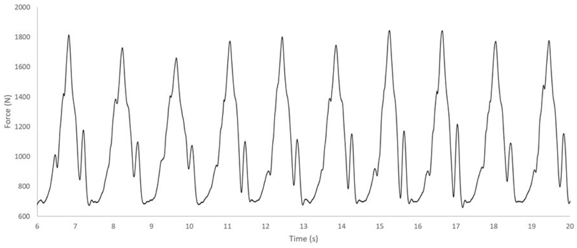


36 kg

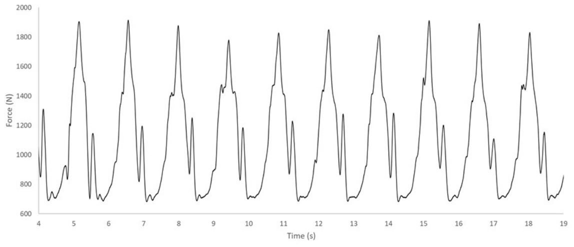

40 kg

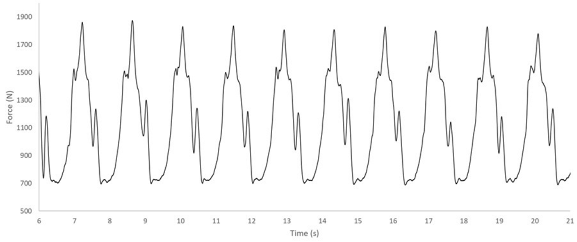


44 kg

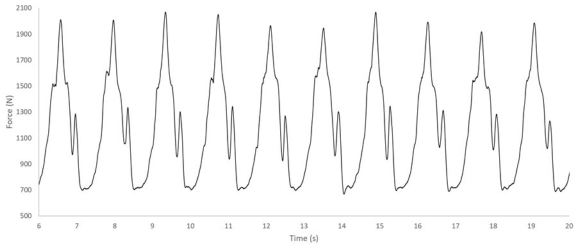


48 kg

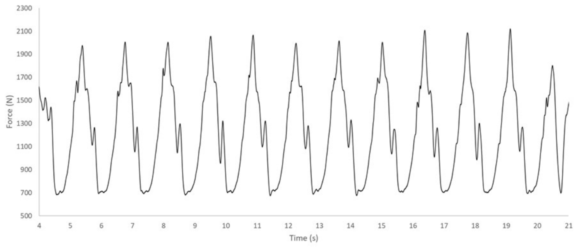


56 kg

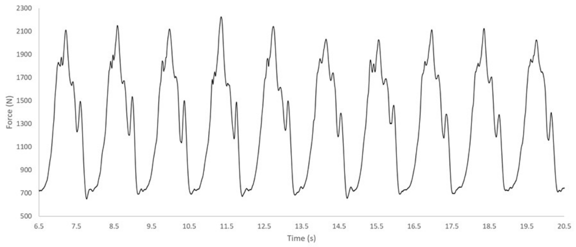


68 kg

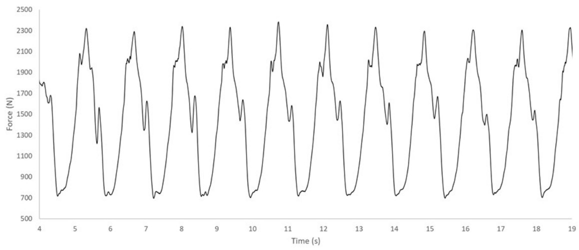
